# Genome-wide association study identifies 30 Loci Associated with Bipolar Disorder

**DOI:** 10.1101/173062

**Authors:** Eli A Stahl, Gerome Breen, Andreas J Forstner, Andrew McQuillin, Stephan Ripke, Vassily Trubetskoy, Manuel Mattheisen, Yunpeng Wang, Jonathan R I Coleman, Héléna A Gaspar, Christiaan A de Leeuw, Stacy Steinberg, Jennifer M Whitehead Pavlides, Maciej Trzaskowski, Tune H Pers, Peter A Holmans, Liam Abbott, Esben Agerbo, Huda Akil, Diego Albani, Ney Alliey-Rodriguez, Thomas D Als, Adebayo Anjorin, Verneri Antilla, Swapnil Awasthi, Judith A Badner, Marie Bækvad-Hansen, Jack D Barchas, Nicholas Bass, Michael Bauer, Richard Belliveau, Sarah E Bergen, Carsten Bøcker Pedersen, Erlend Bøen, Marco Boks, James Boocock, Monika Budde, William Bunney, Margit Burmeister, Jonas Bybjerg-Grauholm, William Byerley, Miquel Casas, Felecia Cerrato, Pablo Cervantes, Kimberly Chambert, Alexander W Charney, Danfeng Chen, Claire Churchhouse, Toni-Kim Clarke, William Coryell, David W Craig, Cristiana Cruceanu, David Curtis, Piotr M Czerski, Anders M Dale, Simone de Jong, Franziska Degenhardt, Jurgen Del-Favero, J Raymond DePaulo, Srdjan Djurovic, Amanda L Dobbyn, Ashley Dumont, Torbjørn Elvsåshagen, Valentina Escott-Price, Chun Chieh Fan, Sascha B Fischer, Matthew Flickinger, Tatiana M Foroud, Liz Forty, Josef Frank, Christine Fraser, Nelson B Freimer, Louise Frisén, Katrin Gade, Diane Gage, Julie Garnham, Claudia Giambartolomei, Marianne Giørtz Pedersen, Jaqueline Goldstein, Scott D Gordon, Katherine Gordon-Smith, Elaine K Green, Melissa J Green, Tiffany A Greenwood, Jakob Grove, Weihua Guan, JoséGuzman Parra, Marian L Hamshere, Martin Hautzinger, Urs Heilbronner, Stefan Herms, Maria Hipolito, Per Hoffmann, Dominic Holland, Laura Huckins, Stéphane Jamain, Jessica S Johnson, Anders Juréus, Radhika Kandaswamy, Robert Karlsson, James L Kennedy, Sarah Kittel-Schneider, Sarah V Knott, James A Knowles, Manolis Kogevinas, Anna C Koller, Ralph Kupka, Catharina Lavebratt, Jacob Lawrence, William B Lawson, Markus Leber, Phil H Lee, Shawn E Levy, Jun Z Li, Chunyu Liu, Susanne Lucae, Anna Maaser, Donald J MacIntyre, Pamela B Mahon, Wolfgang Maier, Lina Martinsson, Steve McCarroll, Peter McGuffin, Melvin G McInnis, James D McKay, Helena Medeiros, Sarah E Medland, Fan Meng, Lili Milani, Grant W Montgomery, Derek W Morris, Thomas W Mühleisen, Niamh Mullins, Hoang Nguyen, Caroline M Nievergelt, Annelie Nordin Adolfsson, Evaristus A Nwulia, Claire O’Donovan, Loes M Olde Loohuis, Anil P S Ori, Lilijana Oruc, Urban Ösby, Roy H Perlis, Amy Perry, Andrea Pfennig, James B Potash, Shaun M Purcell, Eline J Regeer, Andreas Reif, Céline S Reinbold, John P Rice, Fabio Rivas, Margarita Rivera, Panos Roussos, Douglas M Ruderfer, Euijung Ryu, Cristina Sánchez-Mora, Alan F Schatzberg, William A Scheftner, Nicholas J Schork, Cynthia Shannon Weickert, Tatyana Shehktman, Paul D Shilling, Engilbert Sigurdsson, Claire Slaney, Olav B Smeland, Janet L Sobell, Christine Søholm Hansen, Anne T Spijker, David St Clair, Michael Steffens, John S Strauss, Fabian Streit, Jana Strohmaier, Szabolcs Szelinger, Robert C Thompson, Thorgeir E Thorgeirsson, Jens Treutlein, Helmut Vedder, Weiqing Wang, Stanley J Watson, Thomas W Weickert, Stephanie H Witt, Simon Xi, Wei Xu, Allan H Young, Peter Zandi, Peng Zhang, Sebastian Zollner, Rolf Adolfsson, Ingrid Agartz, Martin Alda, Lena Backlund, Bernhard T Baune, Frank Bellivier, Wade H Berrettini, Joanna M Biernacka, Douglas H R Blackwood, Michael Boehnke, Anders D Børglum, Aiden Corvin, Nicholas Craddock, Mark J Daly, Udo Dannlowski, Tõnu Esko, Bruno Etain, Mark Frye, Janice M Fullerton, Elliot S Gershon, Michael Gill, Fernando Goes, Maria Grigoroiu-Serbanescu, Joanna Hauser, David M Hougaard, Christina M Hultman, Ian Jones, Lisa A Jones, RenéS Kahn, George Kirov, Mikael Landén, Marion Leboyer, Cathryn M Lewis, Qingqin S Li, Jolanta Lissowska, Nicholas G Martin, Fermin Mayoral, Susan L McElroy, Andrew M McIntosh, Francis J McMahon, Ingrid Melle, Andres Metspalu, Philip B Mitchell, Gunnar Morken, Ole Mors, Preben Bo Mortensen, Bertram Müller-Myhsok, Richard M Myers, Benjamin M Neale, Vishwajit Nimgaonkar, Merete Nordentoft, Markus M Nöthen, Michael C O’Donovan, Ketil J Oedegaard, Michael J Owen, Sara A Paciga, Carlos Pato, Michele T Pato, Danielle Posthuma, Josep Antoni Ramos-Quiroga, Marta Ribasés, Marcella Rietschel, Guy A Rouleau, Martin Schalling, Peter R Schofield, Thomas G Schulze, Alessandro Serretti, Jordan W Smoller, Hreinn Stefansson, Kari Stefansson, Eystein Stordal, Patrick F Sullivan, Gustavo Turecki, Arne E Vaaler, Eduard Vieta, John B Vincent, Thomas Werge, John I Nurnberger, Naomi R Wray, Arianna Di Florio, Howard J Edenberg, Sven Cichon, Roel A Ophoff, Laura J Scott, Ole A Andreassen, John Kelsoe, Pamela Sklar

**Affiliations:** Department of Genetics and Genomic Sciences, Icahn School of Medicine at Mount Sinai, New York, NY, US; Department of Psychiatry, Icahn School of Medicine at Mount Sinai, New York, NY, US; Medical and Population Genetics, Broad Institute, Cambridge, MA, US; MRC Social, Genetic and Developmental Psychiatry Centre, King’s College London, London, GB; NIHR BRC for Mental Health, King’s College London, London, GB; Department of Biomedicine, University of Basel, Basel, CH; Department of Psychiatry (UPK), University of Basel, Basel, CH; Institute of Human Genetics, University of Bonn, Bonn, DE; Life&Brain Center, Department of Genomics, University of Bonn, Bonn, DE; Institute of Medical Genetics and Pathology, University Hospital Basel, Basel, CH; Division of Psychiatry, University College London, London, GB; Stanley Center for Psychiatric Research, Broad Institute, Cambridge, MA, US; Department of Psychiatry and Psychotherapy, Charité - Universitätsmedizin, Berlin, DE; Analytic and Translational Genetics Unit, Massachusetts General Hospital, Boston, MA, US; iSEQ, Center for Integrative Sequencing, Aarhus University, Aarhus, DK; Department of Biomedicine - Human Genetics, Aarhus University, Aarhus, DK; Department of Clinical Neuroscience, Centre for Psychiatry Research, Karolinska Institutet, Stockholm, SE; Department of Psychiatry, Psychosomatics and Psychotherapy, Center of Mental Health, University Hospital Würzburg, Würzburg, DE; iPSYCH, The Lundbeck Foundation Initiative for Integrative Psychiatric Research, DK; Institute of Biological Psychiatry, Mental Health Centre Sct. Hans, Copenhagen, DK; Institute of Clinical Medicine, University of Oslo, Oslo, NO; Department of Complex Trait Genetics, Center for Neurogenomics and Cognitive Research, Amsterdam Neuroscience, Vrije Universiteit Amsterdam, Amsterdam, NL; deCODE Genetics / Amgen, Reykjavik, IS; Queensland Brain Institute, The University of Queensland, Brisbane, QLD, AU; Institute for Molecular Bioscience, The University of Queensland, Brisbane, QLD, AU; Division of Endocrinology and Center for Basic and Translational Obesity Research, Boston Children’s Hospital, Boston, MA, US; Medical Research Council Centre for Neuropsychiatric Genetics and Genomics, Division of Psychological Medicine and Clinical Neurosciences, Cardiff University, Cardiff, GB; National Centre for Register-Based Research, Aarhus University, Aarhus, DK; Centre for Integrated Register-based Research, Aarhus University, Aarhus, DK; Molecular & Behavioral Neuroscience Institute, University of Michigan, Ann Arbor, MI, US; NEUROSCIENCE, Istituto Di Ricerche Farmacologiche Mario Negri, Milano, IT; Department of Psychiatry and Behavioral Neuroscience, University of Chicago, Chicago, IL, US; Psychiatry, Berkshire Healthcare NHS Foundation Trust, Bracknell, GB; Psychiatry, Rush University Medical Center, Chicago, IL, US; Center for Neonatal Screening, Department for Congenital Disorders, Statens Serum Institut, Copenhagen, DK; Department of Psychiatry, Weill Cornell Medical College, New York, NY, US; Department of Psychiatry and Psychotherapy, University Hospital Carl Gustav Carus, Technische Universität Dresden, Dresden, DE; Department of Medical Epidemiology and Biostatistics, Karolinska Institutet, Stockholm, SE; Department of Psychiatric Research, Diakonhjemmet Hospital, Oslo, NO; Psychiatry, UMC Utrecht Hersencentrum Rudolf Magnus, Utrecht, NL; Human Genetics, University of California Los Angeles, Los Angeles, CA, US; Institute of Psychiatric Phenomics and Genomics (IPPG), University Hospital, LMU Munich, Munich, DE; Department of Psychiatry and Human Behavior, University of California, Irvine, Irvine, CA, US; Molecular & Behavioral Neuroscience Institute and Department of Computational Medicine & Bioinformatics, University of Michigan, Ann Arbor, MI, US; Psychiatry, University of California San Francisco, San Francisco, CA, US; Instituto de Salud Carlos III, Biomedical Network Research Centre on Mental Health (CIBERSAM), Madrid, ES; Department of Psychiatry, Hospital Universitari Vall d’Hebron, Barcelona, ES; Department of Psychiatry and Forensic Medicine, Universitat Autònoma de Barcelona, Barcelona, ES; Psychiatric Genetics Unit, Group of Psychiatry Mental Health and Addictions, Vall d’Hebron Research Institut (VHIR), Universitat Autònoma de Barcelona, Barcelona, ES; Department of Psychiatry, Mood Disorders Program, McGill University Health Center, Montreal, QC, CA; Division of Psychiatry, University of Edinburgh, Edinburgh, GB; University of Iowa Hospitals and Clinics, Iowa City, IA, US; Translational Genomics, USC, Phoenix, AZ, US; Department of Translational Research in Psychiatry, Max Planck Institute of Psychiatry, Munich, DE; Centre for Psychiatry, Queen Mary University of London, London, GB; UCL Genetics Institute, University College London, London, GB; Department of Psychiatry, Laboratory of Psychiatric Genetics, Poznan University of Medical Sciences, Poznan, PL; Department of Neurosciences, University of California San Diego, La Jolla, CA, US; Department of Radiology, University of California San Diego, La Jolla, CA, US; Department of Psychiatry, University of California San Diego, La Jolla, CA, US; Department of Cognitive Science, University of California San Diego, La Jolla, CA, US; Applied Molecular Genomics Unit, VIB Department of Molecular Genetics, University of Antwerp, Antwerp, Belgium; Department of Psychiatry and Behavioral Sciences, Johns Hopkins University School of Medicine, Baltimore, MD, US; Department of Medical Genetics, Oslo University Hospital Ullevål, Oslo, NO; NORMENT, KG Jebsen Centre for Psychosis Research, Department of Clinical Science, University of Bergen, Bergen, NO; Department of Neurology, Oslo University Hospital, Oslo, NO; NORMENT, KG Jebsen Centre for Psychosis Research, Oslo University Hospital, Oslo, NO; Center for Statistical Genetics and Department of Biostatistics, University of Michigan, Ann Arbor, MI, US; Department of Medical & Molecular Genetics, Indiana University, Indianapolis, IN, US; Department of Genetic Epidemiology in Psychiatry, Central Institute of Mental Health, Medical Faculty Mannheim, Heidelberg University, Mannheim, DE; Center for Neurobehavioral Genetics, University of California Los Angeles, Los Angeles, CA, US; Department of Molecular Medicine and Surgery, Karolinska Institutet and Center for Molecular Medicine, Karolinska University Hospital, Stockholm, SE; Department of Clinical Neuroscience, Karolinska Institutet and Center for Molecular Medicine, Karolinska University Hospital, Stockholm, SE; Child and Adolescent Psychiatry Research Center, Stockholm, SE; Department of Psychiatry and Psychotherapy, University Medical Center Göttingen, Göttingen, DE; Department of Psychiatry, Dalhousie University, Halifax, NS, CA; Genetics and Computational Biology, QIMR Berghofer Medical Research Institute, Brisbane, QLD, AU; Department of Psychological Medicine, University of Worcester, Worcester, GB; School of Biomedical and Healthcare Sciences, Plymouth University Peninsula Schools of Medicine and Dentistry, Plymouth, GB; School of Psychiatry, University of New South Wales, Sydney, NSW, AU; Bioinformatics Research Centre, Aarhus University, Aarhus, DK; Biostatistics, University of Minnesota System, Minneapolis, MN, US; Mental Health Department, University Regional Hospital, Biomedicine Institute (IBIMA), Málaga, ES; Department of Psychology, Eberhard Karls Universität Tübingen, Tubingen, DE; Department of Psychiatry and Behavioral Sciences, Howard University Hospital, Washington, DC, US; Center for Multimodal Imaging and Genetics, University of California San Diego, La Jolla, CA, US; Psychiatrie Translationnelle, Inserm U955, Créteil, FR; Faculté de Médecine, Université Paris Est, Créteil, FR; Campbell Family Mental Health Research Institute, Centre for Addiction and Mental Health, Toronto, ON, CA; Neurogenetics Section, Centre for Addiction and Mental Health, Toronto, ON, CA; Department of Psychiatry, University of Toronto, Toronto, ON, CA; Institute of Medical Sciences, University of Toronto, Toronto, ON, CA; Department of Psychiatry, Psychosomatic Medicine and Psychotherapy, University Hospital Frankfurt, Frankfurt am Main, DE; Cell Biology, SUNY Downstate Medical Center College of Medicine, Brooklyn, NY, US; Institute for Genomic Health, SUNY Downstate Medical Center College of Medicine, Brooklyn, NY, US; Center for Research in Environmental Epidemiology (CREAL), Barcelona, ES; Psychiatry, Altrecht, Utrecht, NL; Psychiatry, GGZ inGeest, Amsterdam, NL; Psychiatry, VU medisch centrum, Amsterdam, NL; Psychiatry, North East London NHS Foundation Trust, Ilford, GB; Clinic for Psychiatry and Psychotherapy, University Hospital Cologne, Cologne, DE; Psychiatric and Neurodevelopmental Genetics Unit, Massachusetts General Hospital, Boston, MA, US; HudsonAlpha Institute for Biotechnology, Huntsville, AL, US; Department of Human Genetics, University of Michigan, Ann Arbor, MI, US; Psychiatry, University of Illinois at Chicago College of Medicine, Chicago, IL, US; Max Planck Institute of Psychiatry, Munich, DE; Mental Health, NHS 24, Glasgow, GB; Division of Psychiatry, Centre for Clinical Brain Sciences, University of Edinburgh, Edinburgh, GB; Psychiatry, Brigham and Women’s Hospital, Boston, MA, US; Department of Psychiatry and Psychotherapy, University of Bonn, Bonn, DE; Department of Genetics, Harvard Medical School, Boston, MA, US; Department of Psychiatry, University of Michigan, Ann Arbor, MI, US; Genetic Cancer Susceptibility Group, International Agency for Research on Cancer, Lyon, FR; Estonian Genome Center, University of Tartu, Tartu, EE; Discipline of Biochemistry, Neuroimaging and Cognitive Genomics (NICOG) Centre, National University of Ireland, Galway, Galway, IE; Neuropsychiatric Genetics Research Group, Dept of Psychiatry and Trinity Translational Medicine Institute, Trinity College Dublin, Dublin, IE; Institute of Neuroscience and Medicine (INM-1), Research Centre Jülich, Jülich, DE; Research/Psychiatry, Veterans Affairs San Diego Healthcare System, San Diego, CA, US; Department of Clinical Sciences, Psychiatry, Umeå University Medical Faculty, Umeå, SE; Department of Clinical Psychiatry, Psychiatry Clinic, Clinical Center University of Sarajevo, Sarajevo, BA; Department of Neurobiology, Care sciences, and Society, Karolinska Institutet and Center for Molecular Medicine, Karolinska University Hospital, Stockholm, SE; Psychiatry, Harvard Medical School, Boston, MA, US; Division of Clinical Research, Massachusetts General Hospital, Boston, MA, US; Outpatient Clinic for Bipolar Disorder, Altrecht, Utrecht, NL; Department of Psychiatry, Washington University in Saint Louis, Saint Louis, MO, US; Department of Biochemistry and Molecular Biology II, Institute of Neurosciences, Center for Biomedical Research, University of Granada, Granada, ES; Department of Neuroscience, Icahn School of Medicine at Mount Sinai, New York, NY, US; Medicine, Psychiatry, Biomedical Informatics, Vanderbilt University Medical Center, Nashville, TN, US; Department of Health Sciences Research, Mayo Clinic, Rochester, MN, US; Psychiatry and Behavioral Sciences, Stanford University School of Medicine, Stanford, CA, US; Rush University Medical Center, Chicago, IL, US; Scripps Translational Science Institute, La Jolla, CA, US; Neuroscience Research Australia, Sydney, NSW, AU; Faculty of Medicine, Department of Psychiatry, School of Health Sciences, University of Iceland, Reykjavik, IS; Div Mental Health and Addiction, Oslo University Hospital, Oslo, NO; NORMENT, University of Oslo, Oslo, NO; Psychiatry and the Behavioral Sciences, University of Southern California, Los Angeles, CA, US; Mood Disorders, PsyQ, Rotterdam, NL; Institute for Medical Sciences, University of Aberdeen, Aberdeen, UK; Research Division, Federal Institute for Drugs and Medical Devices (BfArM), Bonn, DE; Centre for Addiction and Mental Health, Toronto, ON, CA; Neurogenomics, TGen, Los Angeles, AZ, US; Psychiatry, Psychiatrisches Zentrum Nordbaden, Wiesloch, DE; Computational Sciences Center of Emphasis, Pfizer Global Research and Development, Cambridge, MA, US; Department of Biostatistics, Princess Margaret Cancer Centre, Toronto, ON, CA; Dalla Lana School of Public Health, University of Toronto, Toronto, ON, CA; Psychological Medicine, Institute of Psychiatry, Psychology & Neuroscience, King’s College London, London, GB; Department of Mental Health, Johns Hopkins University Bloomberg School of Public Health, Baltimore, MD, US; Institute of Genetic Medicine, Johns Hopkins University School of Medicine, Baltimore, MD, US; NORMENT, KG Jebsen Centre for Psychosis Research, Division of Mental Health and Addiction, Institute of Clinical Medicine and Diakonhjemmet Hospital, University of Oslo, Oslo, NO; National Institute of Mental Health, Klecany, CZ; Discipline of Psychiatry, University of Adelaide, Adelaide, SA, AU; Department of Psychiatry and Addiction Medicine, Assistance Publique - Hôpitaux de Paris, Paris, FR; Paris Bipolar and TRD Expert Centres, FondaMental Foundation, Paris, FR; UMR-S1144 Team 1: Biomarkers of relapse and therapeutic response in addiction and mood disorders, INSERM, Paris, FR; Psychiatry, Université Paris Diderot, Paris, FR; Psychiatry, University of Pennsylvania, Philadelphia, PA, US; Department of Psychiatry, University of Münster, Münster, DE; Division of Endocrinology, Children’s Hospital Boston, Boston, MA, US; Centre for Affective Disorders, Institute of Psychiatry, Psychology and Neuroscience, London, GB; Department of Psychiatry & Psychology, Mayo Clinic, Rochester, MN, US; School of Medical Sciences, University of New South Wales, Sydney, NSW, AU; Department of Human Genetics, University of Chicago, Chicago, IL, US; Biometric Psychiatric Genetics Research Unit, Alexandru Obregia Clinical Psychiatric Hospital, Bucharest, RO; Institute of Neuroscience and Physiology, University of Gothenburg, Gothenburg, SE; INSERM, Paris, FR; Department of Medical & Molecular Genetics, King’s College London, London, GB; Neuroscience Therapeutic Area, Janssen Research and Development, LLC, Titusville, NJ, US; Cancer Epidemiology and Prevention, M. Sklodowska-Curie Cancer Center and Institute of Oncology, Warsaw, PL; School of Psychology, The University of Queensland, Brisbane, QLD, AU; Research Institute, Lindner Center of HOPE, Mason, OH, US; Centre for Cognitive Ageing and Cognitive Epidemiology, University of Edinburgh, Edinburgh, GB; Human Genetics Branch, Intramural Research Program, National Institute of Mental Health, Bethesda, MD, US; Division of Mental Health and Addiction, Oslo University Hospital, Oslo, NO; Division of Mental Health and Addiction, University of Oslo, Institute of Clinical Medicine, Oslo, NO; Institute of Molecular and Cell Biology, University of Tartu, Tartu, EE; Mental Health, Faculty of Medicine and Health Sciences, Norwegian University of Science and Technology - NTNU, Trondheim, NO; Psychiatry, St Olavs University Hospital, Trondheim, NO; Psychosis Research Unit, Aarhus University Hospital, Risskov, DK; Munich Cluster for Systems Neurology (SyNergy), Munich, DE; University of Liverpool, Liverpool, GB; Psychiatry and Human Genetics, University of Pittsburgh, Pittsburgh, PA, US; Mental Health Services in the Capital Region of Denmark, Mental Health Center Copenhagen, University of Copenhagen, Copenhagen, DK; Division of Psychiatry, Haukeland Universitetssjukehus, Bergen, NO; Faculty of Medicine and Dentistry, University of Bergen, Bergen, NO; Human Genetics and Computational Biomedicine, Pfizer Global Research and Development, Groton, CT, US; College of Medicine Institute for Genomic Health, SUNY Downstate Medical Center College of Medicine, Brooklyn, NY, US; Department of Clinical Genetics, Amsterdam Neuroscience, Vrije Universiteit Medical Center, Amsterdam, NL; Department of Neurology and Neurosurgery, McGill University, Faculty of Medicine, Montreal, QC, CA; Montreal Neurological Institute and Hospital, Montreal, QC, CA; Department of Biomedical and NeuroMotor Sciences, University of Bologna, Bologna, IT; Department of Psychiatry, Massachusetts General Hospital, Boston, MA, US; Psychiatric and Neurodevelopmental Genetics Unit (PNGU), Massachusetts General Hospital, Boston, MA, US; Faculty of Medicine, University of Iceland, Reykjavik, IS; Department of Psychiatry, Hospital Namsos, Namsos, NO; Department of Neuroscience, Norges Teknisk Naturvitenskapelige Universitet Fakultet for naturvitenskap og teknologi, Trondheim, NO; Department of Genetics, University of North Carolina at Chapel Hill, Chapel Hill, NC, US; Department of Psychiatry, University of North Carolina at Chapel Hill, Chapel Hill, NC, US; Department of Psychiatry, McGill University, Montreal, QC, CA; Dept of Psychiatry, Sankt Olavs Hospital Universitetssykehuset i Trondheim, Trondheim, NO; Clinical Institute of Neuroscience, Hospital Clinic, University of Barcelona, IDIBAPS, CIBERSAM, Barcelona, ES; Institute of Biological Psychiatry, MHC Sct. Hans, Mental Health Services Copenhagen, Roskilde, DK; Department of Clinical Medicine, University of Copenhagen, Copenhagen, DK; Psychiatry, Indiana University School of Medicine, Indianapolis, IN, US; Biochemistry and Molecular Biology, Indiana University School of Medicine, Indianapolis, IN, US

## Abstract

Bipolar disorder is a highly heritable psychiatric disorder that features episodes of mania and depression. We performed the largest genome-wide association study to date, including 20,352 cases and 31,358 controls of European descent, with follow-up analysis of 822 sentinel variants at loci with P<1×10^-4^ in an independent sample of 9,412 cases and 137,760 controls. In the combined analysis, 30 loci reached genome-wide significant evidence for association, of which 20 were novel. These significant loci contain genes encoding ion channels and neurotransmitter transporters (*CACNA1C*, *GRIN2A*, *SCN2A*, *SLC4A1*), synaptic components (*RIMS1*, *ANK3*), immune and energy metabolism components. Bipolar disorder type I (depressive and manic episodes; ^~^73% of our cases) is strongly genetically correlated with schizophrenia whereas bipolar disorder type II (depressive and hypomanic episodes; ^~^17% of our cases) is more strongly correlated with major depressive disorder. These findings address key clinical questions and provide potential new biological mechanisms for bipolar disorder.

## INTRODUCTION

Bipolar disorder (BD) is a severe neuropsychiatric disorder characterized by recurrent episodes of mania and depression which affect thought, perception, emotion, and social behaviour. A lifetime prevalence of 1-2%, elevated morbidity and mortality, onset in young adulthood, and a frequently chronic course make BD a major public health problem and a leading cause of the global burden of disease ^1^. Clinical, twin and molecular genetic data all strongly suggest that BD is a multifactorial disorder ^2^. Based on twin studies, the overall heritability of BD has been estimated to be more than 70% ^3,4^, suggesting a substantial involvement of genetic factors in the development of the disorder, although non-genetic factors also influence risk.

BD can be divided into two main clinical subtypes ^5,6^: bipolar I disorder (BD1) and bipolar II disorder (BD2). In BD1, manic episodes typically alternate with depressive episodes during the course of illness. Diagnosis of BD2 is based on the lifetime occurrence of at least one depressive and one hypomanic (but no manic) episode. Although modern diagnostic systems retain the Kraepelinian dichotomy ^7^ between BD and schizophrenia, the distinction between the two disorders is not always clear-cut, and patients who display clinical features of both disorders may receive a diagnosis of schizoaffective disorder (SAB). Likewise, in genetic studies the two diagnoses are usually treated separately, although recent epidemiological and molecular genetic studies provide strong evidence for some overlap between the genetic contributions to their etiology ^2,8^.

Recent genome-wide association studies (GWAS) in BD have identified a number of significant associations between disease status and common genetic variants ^9–23^. The first large collaborative BD GWAS by the multinational Psychiatric Genomics Consortium (PGC) Bipolar Disorder Working Group comprised 7,481 BD patients and 9,250 controls and identified four genome-wide significant loci ^9^. Three subsequent meta-analyses that included the PGC BD data ^10,12,18^ identified an additional 5 loci.

Estimates of the proportion of variance in liability attributable to common variants genome-wide (SNP-heritability) indicate that ^~^30% of the heritability for BD is due to common genetic variants ^8^. To date, only a small fraction of this heritability is explained by associated loci, but results from other human complex traits suggest that many more will be identified by increasing the sample size of GWAS ^24^. Here, we report the second GWAS of the PGC Bipolar Disorder Working Group, comprising 20,352 cases and 31,358 controls of European descent in a single, systematic analysis, with follow up of top findings in an independent sample of 9,412 cases and 137,760 controls. Some of our findings reinforce specific hypotheses regarding BD neurobiology; however, the majority of the findings suggest new biological insights.

## RESULTS

### GWAS of bipolar disorder (BD)

We performed a GWAS meta-analysis of 32 cohorts from 14 countries in Europe, North America and Australia (**Supplementary Table 1A**), totaling 20,352 cases and 31,358 controls of European descent (effective sample size 46,582). This is the largest GWAS of BD to date and includes 6,328 case and 7,963 control samples not previously reported, a 2.7-fold increase in the number of cases compared to our previous GWAS ^9^. We imputed variant dosages using the 1,000 Genomes reference panel (see Methods), retaining association results for 9,372,253 autosomal variants with imputation quality score INFO > 0.3 and minor allele frequency ≥ 1% in both cases and controls. We performed logistic regression of case status on imputed variant dosage using genetic ancestry covariates. The resulting genomic inflation factor (λ_GC_) was 1.23 and scaled to 1,000 cases and 1,000 controls (λ_1000_) was 1.01 (**Supplementary Figure 1**). The LD-score regression intercept did not significantly differ from one, indicating that the observed genomic inflation is indicative of polygenicity rather than stratification or cryptic population structure ^25^. The LD-score regression SNP-heritability estimates for BD were 0.17-0.23 (on the liability scale, assuming population lifetime risk of 0.5-2%). See **Supplementary Table 1A**, **Online Methods** and **Supplementary Note** for sample and method details.

We find a marked increase in phenotypic variance explained by genomewide polygenic risk scores (PRS) compared to previous publications (sample size weighted mean observed Nagelkerke’s R^2^ = 0.08 across datasets, liability scale R^2^=0.04, for P-threshold 0.01; **Supplementary Figure 2** and **Supplementary Table 2**). Among the different datasets, we observed no association between the PRS and: (i) the gender distribution of the BD cases (p=0.51); (ii) the proportion of cases with psychosis (p=0.61); (iii) the proportion with a family history of BD (p=0.82); or (iv) the median age of onset for BD (p=0.64). In our primary genome-wide analysis, we identified 19 loci exceeding genome-wide significance (P< 5×10^-8^).

### Follow-up of suggestive loci in additional samples

We meta-analyzed lead variants that were significant at P<1×10^-4^ in our discovery meta-analysis, (a total of 794 autosomal and 28 X chromosome variants) with follow-up samples totaling 9,412 cases and 137,760 controls of European ancestry (**Supplementary Note** and **Supplementary Table 1B**). Thirty autosomal loci achieved combined sample genome-wide significance (P< 5×10^-8^) (**Figure 1**, **Table 1**, **Supplementary Figure 3**, **Supplementary Table 3**). These include 19 loci that were significant only in the combined analysis, of which three were reported to have genome-wide significant SNPs in previous studies (*ADCY2* ^18^, *POU3F2* ^18^, *ANK3* ^12,18^), and 11 that were significant in our GWAS. Eight variants were genome-wide significant in the GWAS but not in the combined analysis. Using effect sizes corrected for winner’s curse ^26,27^ for each of the 19 variants with GWAS P<5×10^-8^, we found that 11 variants achieving genome-wide significance in our combined analysis is within the expected range (Poisson binomial test P = 0.29, **Supplementary Note** and **Supplementary Figure 4**).

**Figure 1.**
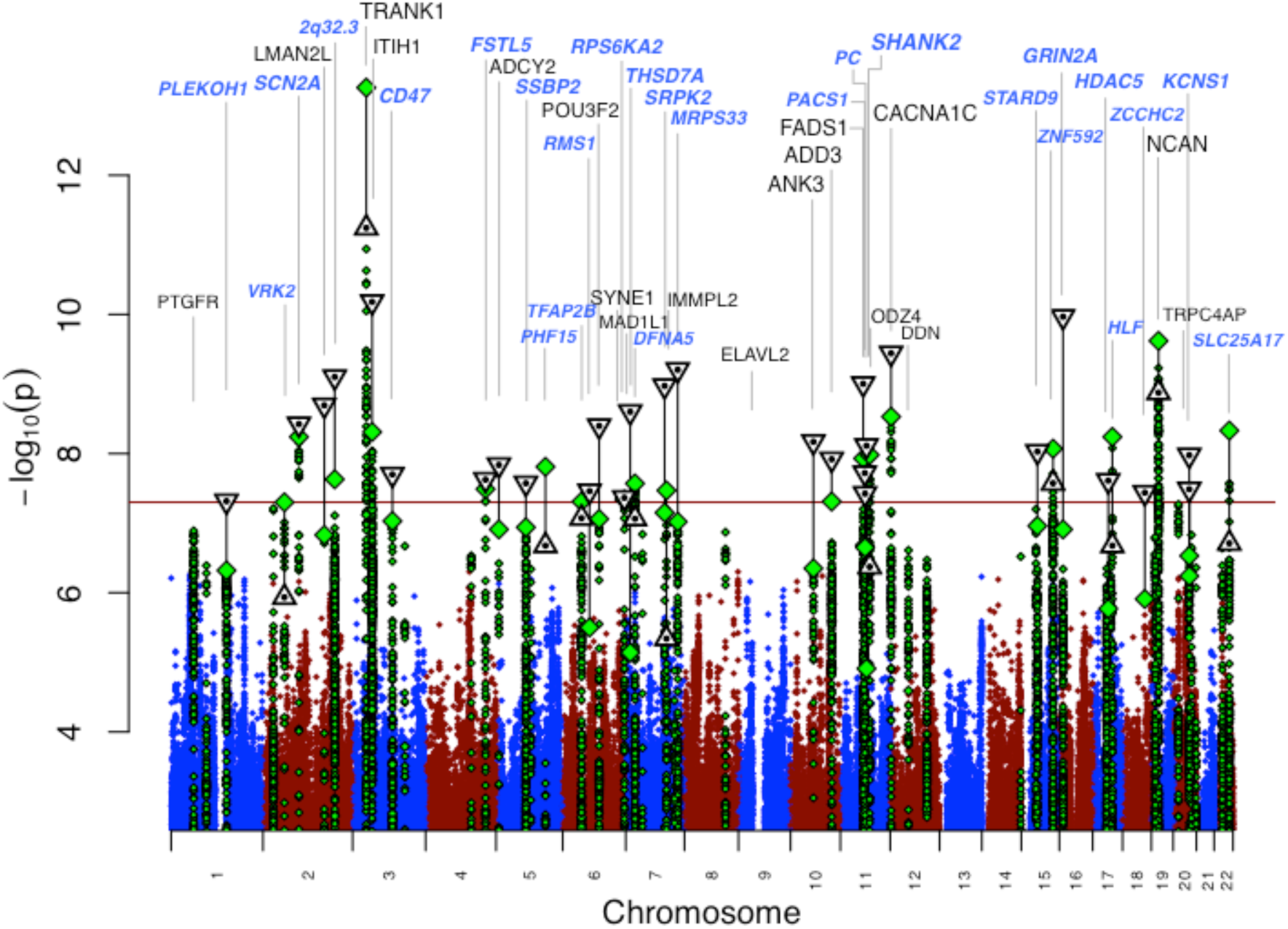
Manhattan plot for our primary genomewide association analysis of 20,352 cases and 31,358 controls. GWAS-log_10_P-values are plotted for all SNPs across chromosomes 1-22 (diamonds, green for loci with lead SNP GWAS P < 10^-6^). Combined GWAS+followup-log_10_P-values for lead SNPs reaching genome-wide significance in either GWAS or combined analysis (triangles, inverted if GWAS+followup-log_10_P > GWAS-log_10_P). Labels correspond to gene symbols previously reported for published loci (black) and the nearest genes for novel loci (blue), at top if GWAS+followup P < 5×10^-8^.

**Table 1.**
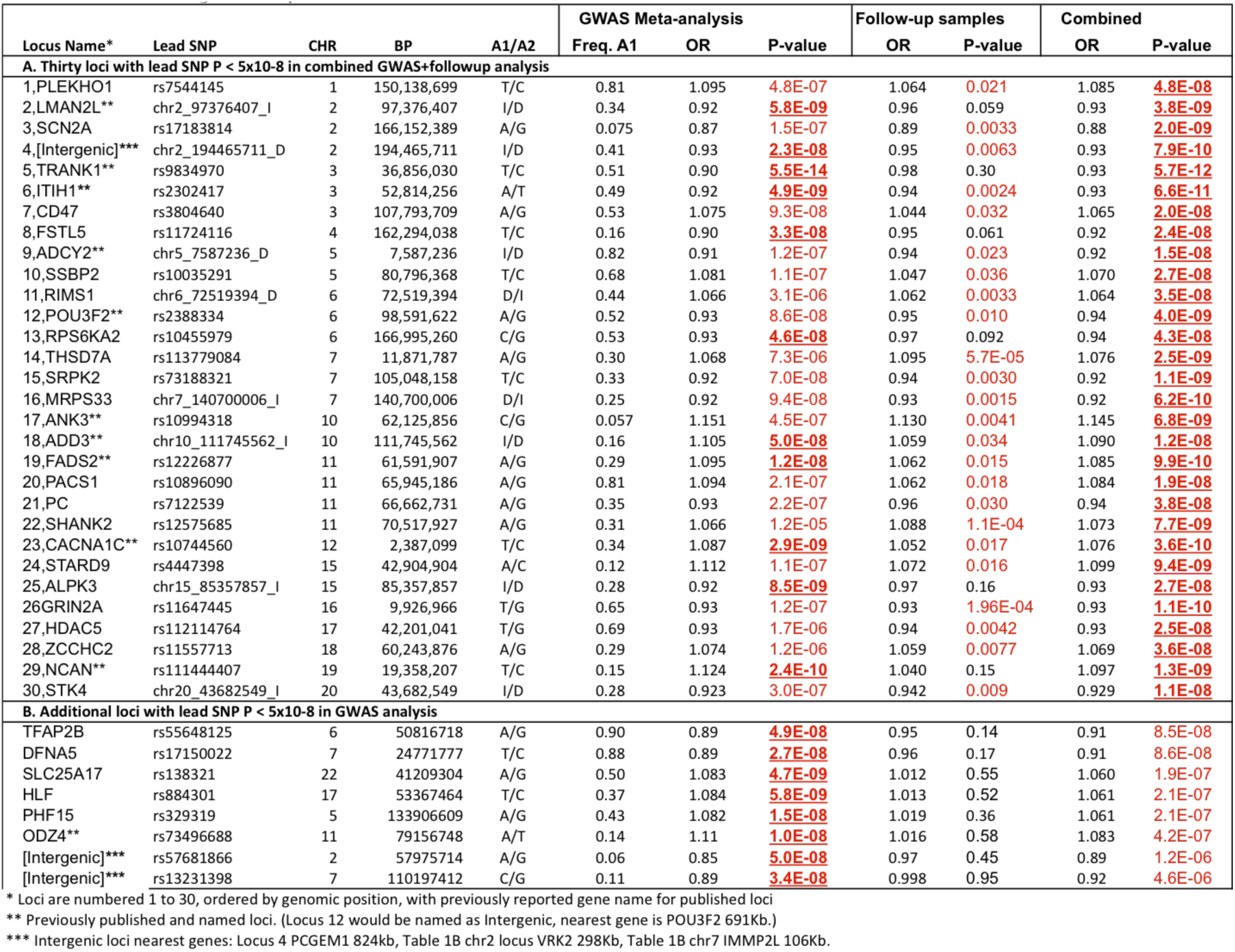
Genome-wide significant bipolar disorder risk loci

Lead variants for the 30 loci achieving genome-wide significance in the combined analysis are shown in **Table 1A**. We show results in **Table 1B** for 8 additional loci with P < 5×10^-8^ in our discovery GWAS but not in the combined analysis. Results for all variants tested in the follow-up study are presented in **Supplementary Table 3**. We refer to loci by the gene name attributed in previous BD GWAS publications, or by the name of the closest gene for novel loci, without implication that the named gene is causal. Of the 30 genome-wide significant loci from our combined analysis, 20 are novel BD risk loci. In **Supplementary Table 4**, we present detailed descriptions of the associated loci and genes, with bioinformatic and literature evidence for their potential roles in BD.

We next asked if the variants tested in the follow-up samples were, in aggregate, consistent with the presence of additional sub genome-wide significant BD association signals. After excluding 47 variants that were genome-wide significant in our GWAS, our combined analysis or previous BD GWAS, 775 variants remained in our follow-up experiment. 551 variants had the same direction of effect in the discovery GWAS and follow-up samples (71% compared to a null expectation of 50%, sign test P < 2.2×10^-16^), and 110 variants had the same direction of effect and were nominally significant (p<0.05) in the follow-up samples (14% compared to an expected value of 2.5%, binomial test P < 2.2×10^-16^). This consistency between our GWAS and follow-up samples suggests that many true BD associations exist among these variants.

To identify additional independent signals, we conducted conditional analyses across each of the 30 significant BD loci (**Supplementary Table 5**). We used the effective number of independent variants based on LD structure within loci ^28^ to calculate a multiple test-corrected significance threshold (P=1.01×10^-5^, see **Supplementary Note**). One locus showed evidence for an independent association signal (rs114534140 in locus #8, *FSTL5;* P_conditional_ = 2×10^-6^). At one locus (#30, STK4 on chr 20), we found two SNPs with genome-wide significance in low LD (r^2^ < 0.1); however, conditional analysis showed that their associations were not independent. Thus only the *FSTL5* locus demonstrated clear evidence of more than one independent association.

### Shared loci and genetic correlations with schizophrenia, depression and other GWAS traits

We next examined the genetic relationships of BD to other psychiatric disorders and traits. Of the 30 genome-wide significant BD loci, 8 also harbor schizophrenia (SCZ) associations ^29-31^. Based on conditional analyses the BD and SCZ associations appear to be independent at 3 of the 8 shared loci (*NCAN*, *TRANK1* and chr7q22.3:105Mb loci) (**Supplementary Table 6**). No genome-wide significant BD locus overlapped with those identified for major depression (DEPR), including 44 risk loci identified in the most recent PGC study based on 130,664 depression cases and 330,470 controls^32^, and those reported in a large study of depressive symptoms or subjective well-being ^33^. As previously reported ^34^, we found substantial and highly significant genetic correlations between BD and SCZ (LD-score regression estimated genetic correlation r_g_ = 0.70, se = 0.020) and between BD and DEPR (r_g_ = 0.35, se = 0.026) The BD and DEPR genetic correlation was similar to that observed for SCZ and DEPR (r_g_ = 0.34, se = 0.025) (**Supplementary Table 7A**).

We found significant genetic correlations between BD and other psychiatric-relevant traits (**Supplementary Table 7B**), including with autism spectrum disorder ^8^ (r_g_ = 0.18, P=2×10^-4^), anorexia nervosa ^35^ (r_g_ = 0.23, P=9×10^-8^), and subjective well-being ^33^ (r_g_ = −0.22, P=4×10^-7^). There was suggestive positive overlap with anxiety disorders (r_g_=0.21, P=0.04)^36^ and neuroticism (r_g_=0.12, P=0.002)^37^. Significant r_g_s were seen with measures of education: college attendance ^38^ (r_g_ = 0.21, P=1=×10^-7^) and education years ^39^ (r_g_=0.20, P=6×10^-14^), but not with childhood IQ ^40^ (r_g_=0.05, P=0.5) or intelligence ^41^ (r_g_=-0.05, P=0.08). Among a large number of BD risk locus SNPs associated with additional traits from GWAS catalog, we found a handful of loci with non-independent associations (in one overlapping locus each with educational attainment, biliary atresia, bone mineral density, lipid-related biomarkers) (**Supplementary Table 6**). Biliary atresia and lipid-related biomarkers, however, did not show significant genetic correlation with BD (**Supplementary Table 7B**).

### BD subtype GWAS

We performed secondary GWAS focusing on three clinically recognized subtypes of bipolar disorder: BD1 (n=14,879 cases), BD2 (n=3,421 cases), and SAB (n=977 cases) (**Supplementary Note**, **Supplementary Tables 1A & 8, Supplementary Figure 5**). We observed variants in 14 loci with genome-wide significance for BD1, 10 of which were in genome-wide significant loci in the combined BD GWAS analysis. Not surprisingly given the sample overlap, 3 of the 4 remaining loci genome-wide significant for BD1 have P < 10^-6^ in either our GWAS or combined analysis. The remaining locus (MAD1L1, chr7:1.9Mb, GWAS P = 2.4×10^-6^) was recently published in two BD GWAS that included Asian samples ^42,43^. We did not observe genome-wide significant results for the smaller BD2 and SAB analyses. BD1, BD2 and SAB all have significant common variant heritabilities (BD1 h^2^_snp_ = 0.25, se = 0.014, P = 3.2×10^-77^; BD2 h^2^_snp_ = 0.11, se = 0.028, P = 5.8×10^-5^; SAB h^2^_snp_= 0.25, se = 0.10, P = 0.0071). Genetic correlations among BD subtypes show that these represent closely related, yet partially distinct, phenotypes (**Supplementary Table 9)**.

Polygenic risk scores and genetic correlations provide support for a continuum of SCZ-BD1-BD2-DEPR genetic effects, with significantly greater genetic SCZ polygenic risk scores (PRS) in BD1 cases than in BD2 cases (min P=5.6×10^-17^, P threshold = 0.1), and greater DEPR PRS in BD2 cases than in BD1 cases (min P=8.5×10^-10^, P threshold = 0.01) (**Figure 2**, **Supplementary Table 10**). Genetic correlations from LD-score regression support these results; genetic correlations were greater for SCZ with BD1 (r_g_ = 0.71, se = 0.025) than with BD2 (r_g_ = 0.51, se = 0.072), with P_diff_ = 0.0056, and were greater for DEPR with BD2 (r_g_ = 0.69, se = 0.093) than with BD1 (r_g_ = 0.30, se = 0.028), with P_diff_ = 2.9×10^-5^ (**Supplementary Table 9**).

**Figure 2.**
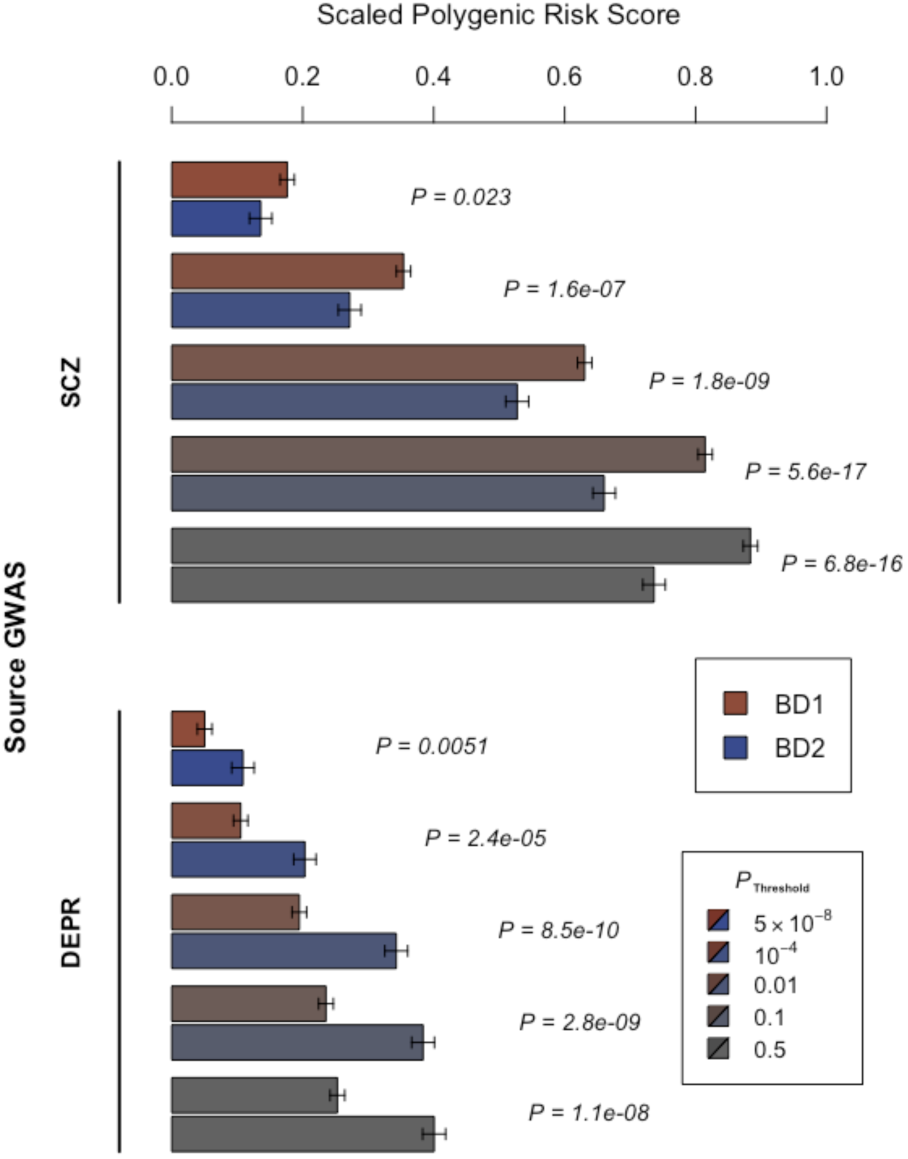
Association of BD1 and BD2 subtypes with schizophrenia (SCZ) and major depression (DEPR) polygenic risk scores (PRS). Shown are mean PRS values (1 s.e. error bars), adjusted for study and ancestry covariates and scaled to the PRS mean and sd in control subjects, in BD1 (red) and BD2 (blue) cases, for increasing source GWAS P-value thresholds (increasing grey) as indicated. P-values (italics) test BD1 vs BD2 mean PRS, in logistic regression of case subtype on PRS with covariates. Results are detailed in Supplementary Table 10.

### Systems biology and *in silico* functional analyses of BD GWAS results

To identify genes with functional variation in gene expression that might explain the associations, we used summary Mendelian randomization (SMR) ^44^ to integrate our BD discovery GWAS with eQTL data from brain dorsolateral prefrontal cortex ^45^ as well as a large-sample whole blood eQTL dataset ^46^ (**Supplemental Table 11**). SMR identified six transcriptome-wide significant genes without signs of heterogeneity between GWAS and eQTL association signals. Among these, four genes were present in four different loci from our combined BD GWAS and follow-up sample meta-analysis: *LMAN2L* (blood), *FADS1* (brain), *NMB* (blood) and *C17ORF65* (blood).

We tested for functional genomic enrichment in our BD GWAS using partitioned LD-score regression ^47^ (**Supplementary Note**, **Supplementary Table 12**). Annotations tested included open chromatin DHS peaks in a range of tissues ^48^, genic annotations, conservation, and a number of functional genomic annotations across tissues. SNP-based BD heritability was most substantially enriched in open chromatin annotations in central nervous system (proportion SNPs = 0.14, proportion h^2^_snp_ = 0.60, enrichment =3.8, P = 4.2 × 10^-17^). We also used DEPICT ^49^ to test for expression of BD associated genes across tissues, and found significant enrichment of central nervous system (P <= 1.3×10^-3^, FDR < 0.01) and neurosecretory system (P <= 2.0×10^-6^, FDR < 0.01) genes (**Supplementary Table 13**).

Finally, we used MAGMA ^50^ to conduct a gene-wise BD GWAS and to test for enrichment of pathways curated from multiple sources (see **Supplementary Note**). We note that significance levels were assigned to genes by physical proximity of SNPs, and do not imply that significant genes are causal for BD. Genic association results included 154 Bonferroni significant genes (MAGMA P_JOINT < 2.8×10^-6^), including 82 genes in 20 genome-wide significant loci, and 73 genes in 27 additional loci that did not reach genome-wide significance in either our GWAS or combined analysis (**Supplementary Table 14**). Nine related pathways were significantly enriched for genes with stronger BD associations (P < 7.0×10^-5^, FDR < 0.05), including abnormal motor coordination/balance pathways (from mice), regulation of insulin secretion and endocannabinoid signaling pathways (**Supplementary Table 15, Supplementary Figure 6**).

## DISCUSSION

We carried out the largest bipolar disorder (BD) GWAS to date and identified 30 genome-wide significant loci, including 20 novel BD risk loci. Previous BD GWAS have reported a total of 20 loci significantly associated with BD^9-23^; twelve of these previously reported loci were not genome-wide significant in our GWAS meta analysis but had P_GWAS_ ≤ 1.3×10^-5^. Of the 19 loci identified in our discovery GWAS, only 11 were genome-wide significant in meta-analysis of our GWAS and follow-up samples. Although these results are not unexpected given small effect sizes and the winner’s curse ^27,51^ (**Supplementary Note** and **Supplementary Figure 4**), genetic heterogeneity has been shown between BD GWAS cohorts^8^. We observed variable polygenic effects between BD subtypes and between cohorts in our study (**Figure 2**, **Supplementary Figure 2**, **Supplementary Tables 2 & 10**) and acknowledge a diversity of clinical case phenotypic criteria among cohorts in our study (**Supplementary Note**). Remarkably, our strongest association signal, observed at the *TRANK1* locus (rs9834970; P_combined_ = 5.7E-12, OR = 0.93), exhibited significant heterogeneity among discovery GWAS cohorts (Cochran’s Q P = 1.9×10^-4^, and did not replicate in the follow-up sample (1-tailed P_followup_ = 0.3) (**Supplementary Figure 3B & 3C**, fifth and first plots respectively). This locus has been observed in recent ^11,12,17,18^ but not earlier BD GWAS ^9,13,20^, surprisingly given its relatively large apparent effect size. Thus, complex polygenic architecture as well as phenotypic heterogeneity among BD GWAS cohorts may contribute to the inconsistency of genome-wide significant findings within and across BD GWAS studies. The observed heterogeneity is a major challenge for GWAS of psychiatric disorders and calls for careful and systematic clinical assessment of cases and controls in addition to continued efforts to collect larger sample sizes.

Of the 30 BD associated loci, 8 also harbor associations ^29-31^ with schizophrenia (SCZ); however, conditional analyses suggest that the BD and SCZ associations at 3 of the 8 shared loci (in the *NCAN*, *TRANK1* and chr7q22.3 [105Mb] loci) may be independent (**Supplementary Table 6**). Differential BD and SCZ associations may represent opportunities to understand the genetic distinctions between these closely related and sometimes clinically difficult to distinguish disorders. We did not find BD loci that overlap with those associated with major depression^32^.

The confirmed association within loci containing *CACNA1C* and other voltage-gated calcium channels supports the rekindled interest in calcium channel antagonists as potential treatments for BD with similar examination ongoing for other genes implicated by current GWAS ^52^. These processes are important in neuronal hyperexcitability^53^, an excess of which has been reported in iPSC derived neurons from BD patients, and which has been shown to be affected by the classic mood stabilizing drug lithium^54^. Other genes within novel associated loci include those coding for neurotransmitter channels (*GRIN2A*), ion channels and transporters (*SCN2A*, *SLC4A1*) and synaptic components (*RIMS1*, *ANK3*). Further study will confirm whether or not these are the causal genes in these loci.

The estimated variance explained by polygenic risk scores (PRS) based on our BD GWAS data is ^~^8% (observed scale; 4% on the liability scale ^55^), an increase from 2.8% from our previous study ^9^. Using PRS, we found that BD1 cases have significantly greater schizophrenia genetic risk than BD2 cases, while BD2 cases have significantly greater major depression genetic risk than BD1 cases, consistent with a spectrum of related psychiatric diagnoses^7,56^. We observe significant positive genetic correlations with educational attainment, but not with either adult or childhood IQ, suggesting that the role of BD genetics in increased educational attainment may be independent of general intelligence. This result is inconsistent with suggestions from epidemiological studies ^57^, but in agreement with a recent clinical study ^58^.

In summary, findings from the largest genome-wide analysis of BD reveal an extensive polygenic genetic architecture of the disease, implicate brain calcium channels and neurotransmitter function in BD etiology, and confirm that BD is part of a spectrum of highly correlated psychiatric and mood disorders.

## ONLINE METHODS

### Methods

#### GWAS and follow-up cohorts

Our discovery GWAS sample was comprised of 32 cohorts from 14 countries in Europe, North America and Australia (**Supplementary Table 1A**), totaling 20,352 cases and 31,358 controls of European descent. A selected set of variants (see below) were tested in 7 follow-up cohorts of European descent (**Supplementary Table 1B**), totalling 9,025 cases and 142,824 controls (N_eff_ = 23,991). The **Supplementary Note** summarizes the source and inclusion/exclusion criteria for cases and controls for each cohort. All cohorts in the initial PGC BD paper were included ^9^. Cases were required to meet international consensus criteria (DSM-IV, ICD-9, or ICD-10) for a lifetime diagnosis of BD established using structured diagnostic instruments from assessments by trained interviewers, clinician-administered checklists, or medical record review. In most cohorts, controls were screened for the absence of lifetime psychiatric disorders and randomly selected from the population.

#### GWAS cohort analysis

We tested 20 principal components for association with BD using logistic regression; seven were significantly associated with phenotype and used in GWAS association analysis (PCs 1-6, 19). In each cohort, we performed logistic regression association tests for BD with imputed marker dosages including 7 principal components to control for population stratification. For all GWAS cohorts, X-chromosome association analyses were conducted separately by sex, and then meta-analyzed across sexes. We also conducted BD1, BD2, and SAB GWAS, retaining only cohorts with at least 35 subtype cases and filtering SNPs for MAF > 0.02. Results were combined across cohorts using an inverse variance-weighted fixed effects meta-analysis ^59^. We used Plink ‘clumping’^60,61^ to identify an LD-pruned set of discovery GWAS meta-analysis BD-associated variants (*P* < 0.0001, and distance >500kb or LD r^2^ < 0.1, n variants =822) for analysis in the follow-up cohorts. Conditional analyses were conducted within each GWAS cohort and meta-analyzed as above.

#### Follow-up cohort analysis

In each follow-up cohort we performed BD association analysis of the 822 selected GWAS variants (when available) including genetic ancestry covariates, following QC and analysis methods of the individual study contributors. We performed inverse variance-weighted fixed-effects meta-analyses of the association results from the follow-up cohorts, and of the discovery GWAS and follow-up analyses.

#### Polygenic risk score (PRS) analyses

We tested PRS for our primary GWAS on each GWAS cohort as a target set, using a GWAS where the target cohort was left out of the meta-analysis (**Supplementary Table 2**). To test genetic overlaps with other psychiatric diseases, we calculated PRS for DEPR and SCZ in our GWAS cohort BD cases ^62^. In pairwise case subtype analyses (**Figure 2**, **Supplementary Table 10**), we regressed subtype case status (BD1 n=8044, BD2 n=3,365, SAB n=977) on the PRS adjusting for ancestry principal components and a cohort indicator using logistic regression, and visualized covariate-adjusted PRS in BD1 and BD2 subtypes (**Figure 2**).

#### Linkage disequilibrium (LD) score regression

LD score regression ^25,63^ was used to conduct SNP-heritability analyses from GWAS summary statistics. LD score regression bivariate genetic correlations attributable to genome-wide common variants were estimated between the full BD GWAS, BD subtype GWASs, and other traits and disorders with LD-Hub ^63^. We also used LD score regression to partition heritability by genomic features ^47^.

#### Relation of BD GWA findings to tissue and cellular gene expression

We used partitioned LD score regression to evaluate which somatic tissues and brain tissues were enriched for BD heritability.^64^ We used summary-data-based Mendelian randomization (SMR) ^44^ to identify loci with strong evidence of causality via gene expression (**Supplementary Table 9**). Since the aim of SMR is to prioritize variants and genes for subsequent studies, a test for heterogeneity excludes regions that may harbor multiple causal loci (pHET < 0.05).

#### Gene-wise and pathway analysis

Guided by rigorous method comparisons conducted by PGC members ^50,65^, p-values quantifying the degree of association of genes and gene sets with BD were generated using MAGMA (v1.06)^50^. We used ENSEMBL gene coordinates for 18,172 genes giving a Bonferroni corrected *P*-value threshold of 2.8×10^-6^. Joint multi-SNP LD-adjusted gene-level p-values were calculated using SNPs 35 kb upstream to 10 kb downstream, adjusting for LD using 1,000 Genomes Project (Phase 3 v5a, MAF ≥ 0.01, European-ancestry subjects)^66^. Gene sets were compiled from multiple sources. Competitive gene set tests were conducted correcting for gene size, variant density, and LD within and between genes. The pathway map (**Supplementary Figure 6**) was constructed using the kernel generative topographic mapping algorithm (k-GTM) as described by ^67^. See **Supplementary Note** for further details.

#### Genome build

All genomic coordinates are given in NCBI Build 37/UCSC hg19.

#### Availability of results

The PGC’s policy is to make genome-wide summary results public. Summary statistics for our meta-analysis of the GWAS cohort samples are available through the PGC (URLs).

## URLs

Psychiatric Genomics Consortium, PGC, https://med.unc.edu/pgc

PGC “ricopili” GWA pipeline, https://github.com/Nealelab/ricopili

1000 Genomes Project multi-ancestry imputation panel, https://mathgen.stats.ox.ac.uk/impute/datadownload1000Gphase1integrated.html

LD-Hub, http://ldsc.broadinstitute.org

GTEx, http://www.gtexportal.org/home/datasets

CommonMind Consortium, http://commonmind.org

## Funding

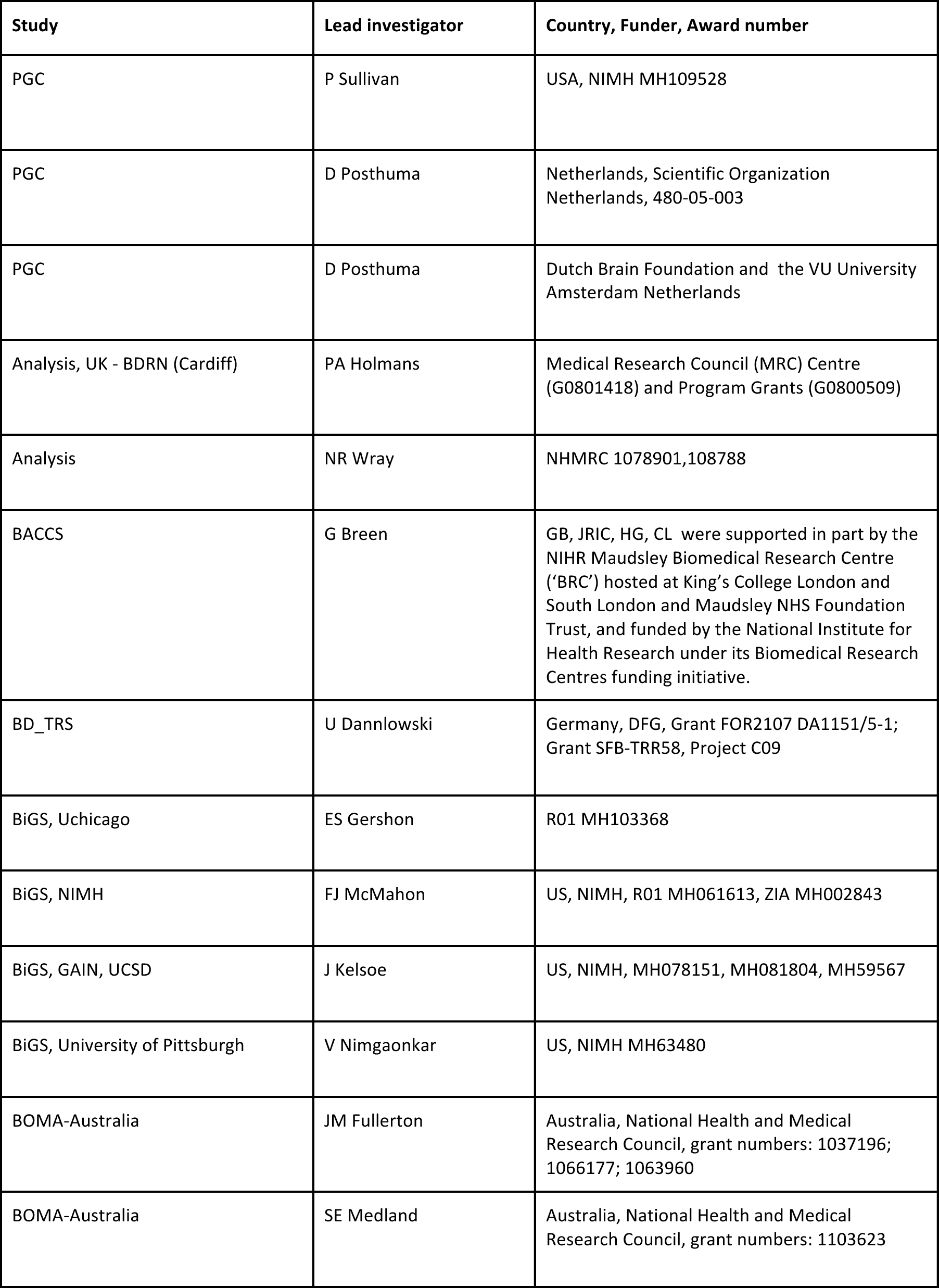
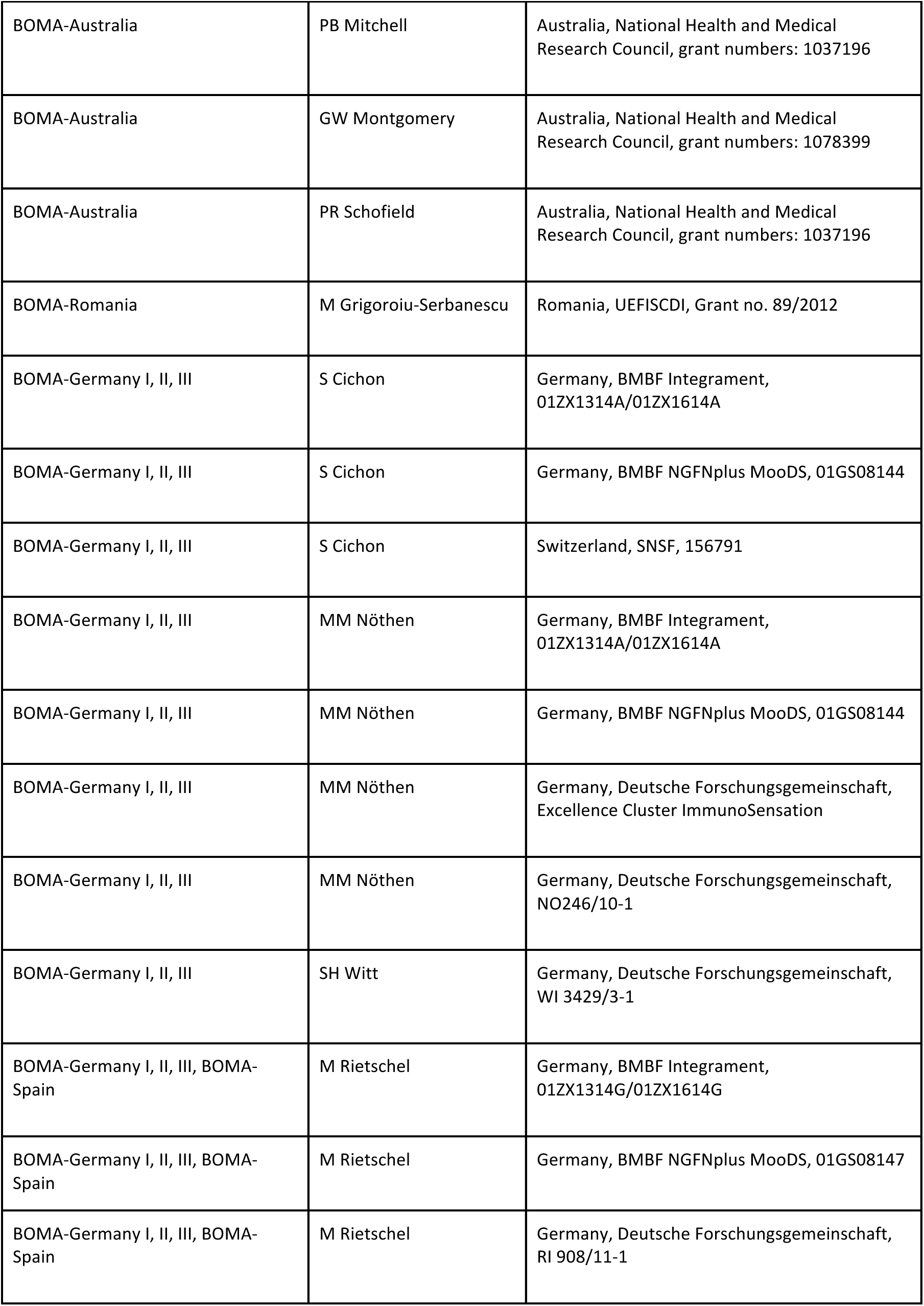
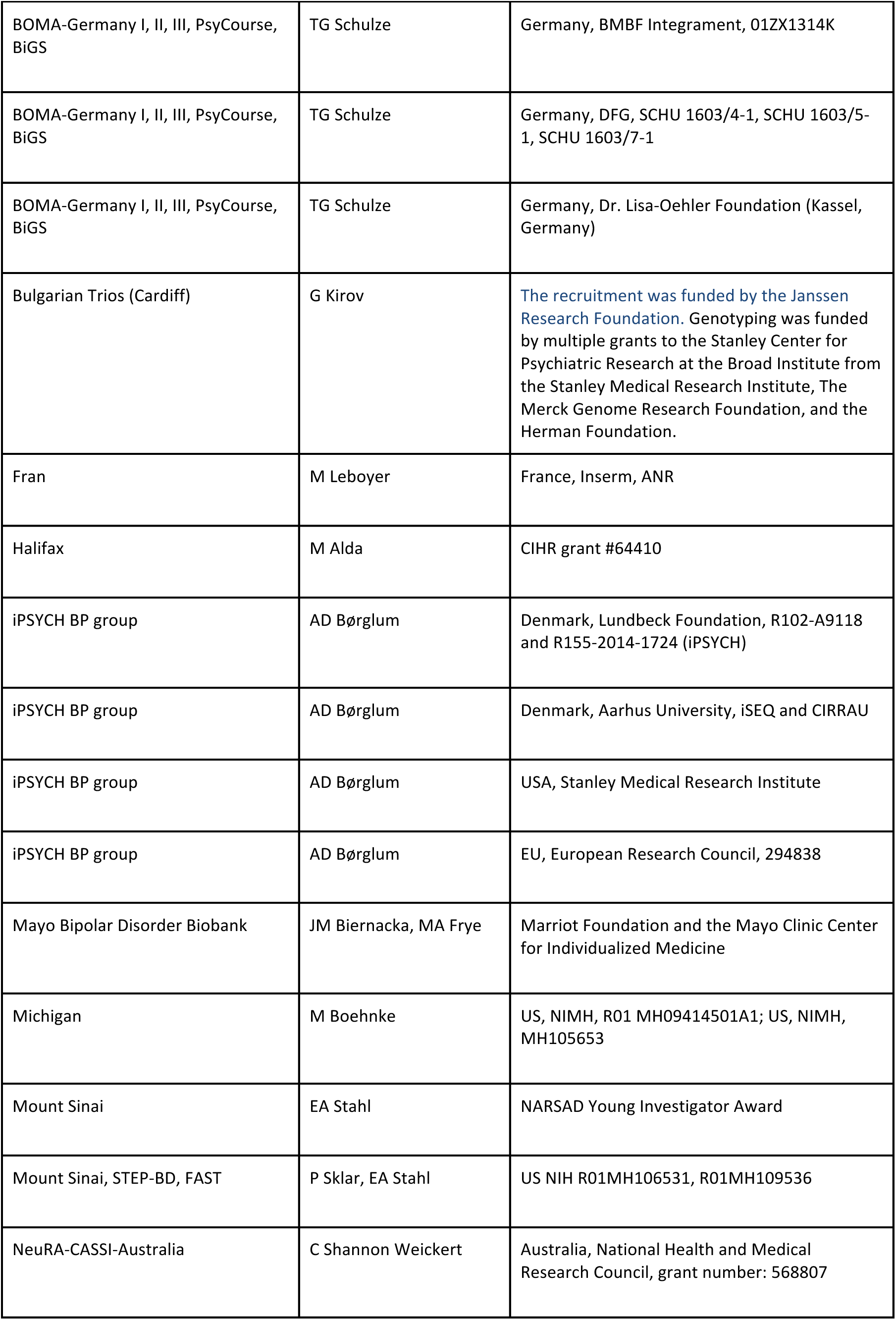
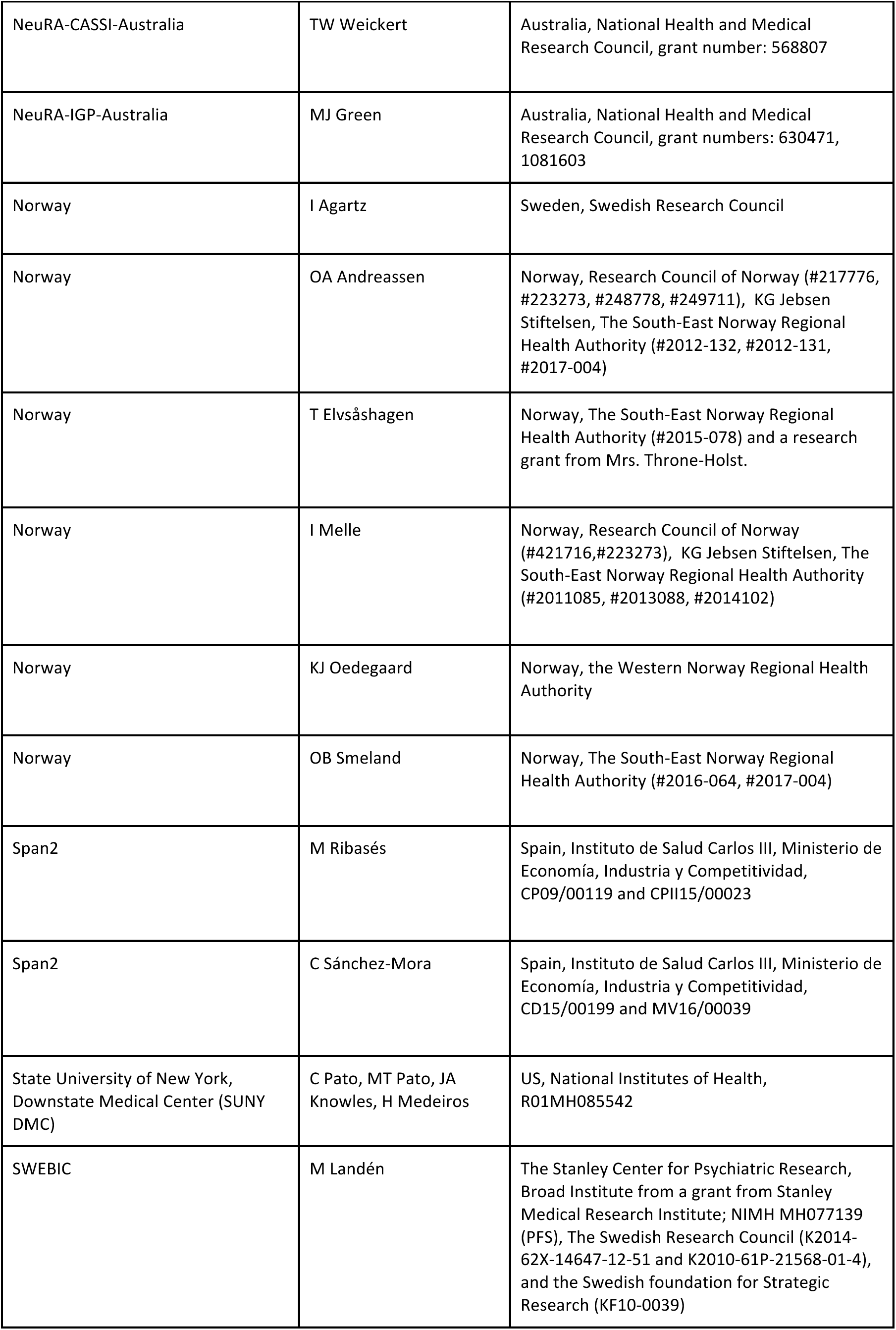
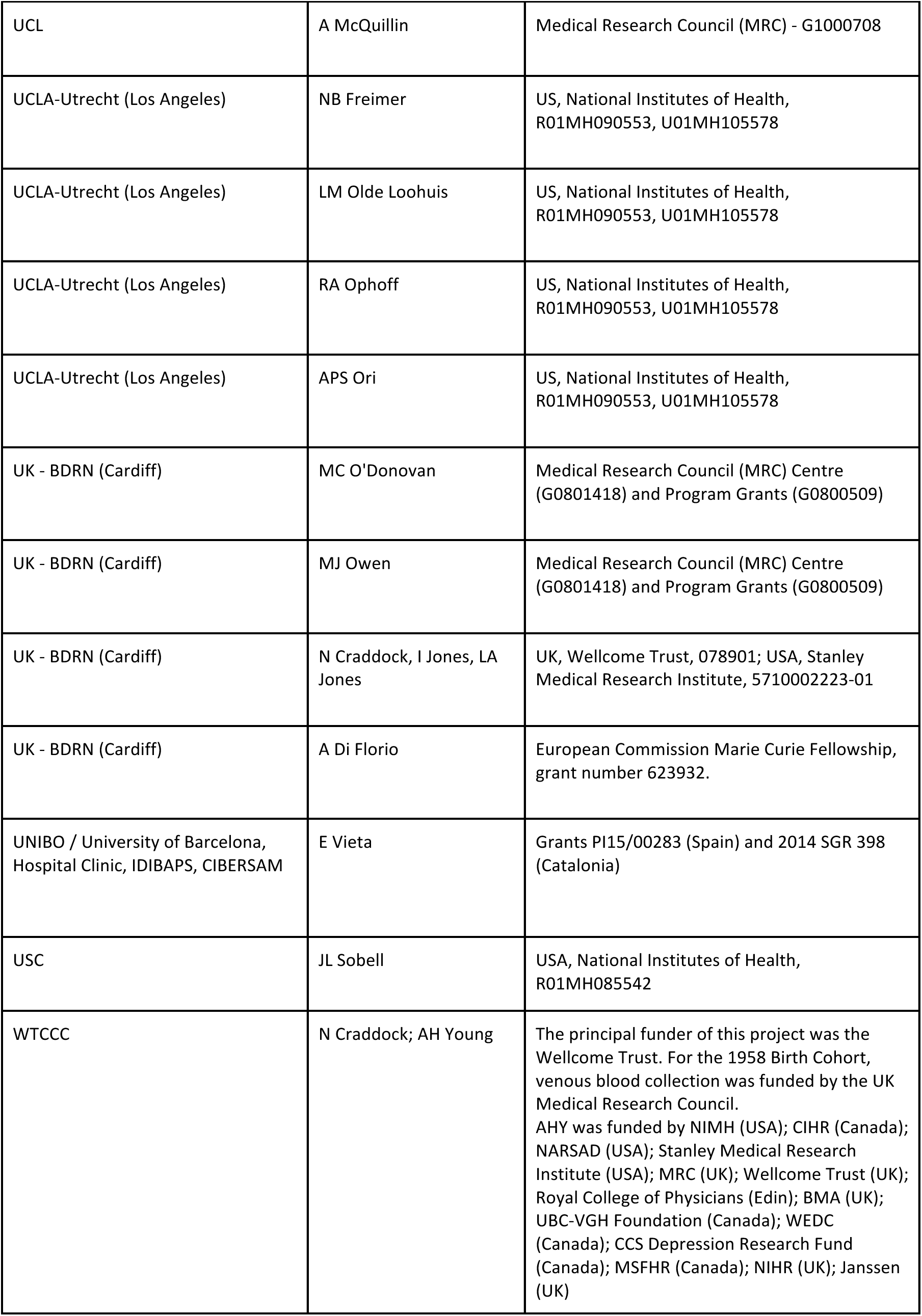

## Acknowledgments

This paper is dedicated to the memory of Psychiatric Genomics Consortium (PGC) founding member and Bipolar disorder working group co-chair Pamela Sklar. We are deeply indebted to the investigators who comprise the PGC, and to the subjects who have shared their life experiences with PGC investigators. The PGC has received major funding from the US National Institute of Mental Health (PGC3: U01 MH109528, PGC2: U01 MH094421, PGC1: U01 MH085520). Statistical analyses were carried out on the NL Genetic Cluster Computer (http://www.geneticcluster.org) hosted by SURFsara.

BACCS: This work was supported in part by the NIHR Maudsley Biomedical Research Centre (‘BRC’) hosted at King’s College London and South London and Maudsley NHS Foundation Trust, and funded by the National Institute for Health Research under its Biomedical Research Centres funding initiative. The views expressed are those of the authors and not necessarily those of the BRC, the NHS, the NIHR or the Department of Health or King’s College London. We gratefully acknowledge capital equipment funding from the Maudsley Charity (Grant Reference 980) and Guy’s and St Thomas’s Charity (Grant Reference STR130505).

BD_TRS: This work was funded by the German Research Foundation (DFG, grant FOR2107 DA1151/5-1 to UD; SFB-TRR58, Project C09 to UD) and the Interdisciplinary Center for Clinical Research (IZKF) of the medical faculty of Münster (grant Dan3/012/17 to UD).

BiGS, GAIN: FJM was supported by the NIMH Intramural Research Program, NIH, DHHS.

BOMA-Australia: JMF would like to thank Janette M O’Neil and Betty C Lynch for their support.

BOMA-Germany I, BOMA-Germany II, BOMA-Germany III, PsyCourse: This work was supported by the German Ministry for Education and Research (BMBF) through the Integrated Network IntegraMent (Integrated Understanding of Causes and Mechanisms in Mental Disorders), under the auspices of the e:Med program (grant 01ZX1314A/01ZX1614A to MMN and SC, grant 01ZX1314G/01ZX1614G to MR, grant 01ZX1314K to TGS). This work was supported by the German Ministry for Education and Research (BMBF) grants NGFNplus MooDS (Systematic Investigation of the Molecular Causes of Major Mood Disorders and Schizophrenia; grant 01GS08144 to MMN and SC, grant 01GS08147 to MR). This work was also supported by the Deutsche Forschungsgemeinschaft (DFG), grant NO246/10-1 to MMN (FOR 2107), grant RI 908/11-1 to MR (FOR 2107), grant WI 3429/3-1 to SHW, grants SCHU 1603/4-1, SCHU 1603/5-1 (KFO 241) and SCHU 1603/7-1 (PsyCourse) to TGS. This work was supported by the Swiss National Science Foundation (SNSF, grant 156791 to SC). MMN is supported through the Excellence Cluster ImmunoSensation. TGS is supported by an unrestricted grant from the Dr. Lisa-Oehler Foundation. AJF received support from the BONFOR Programme of the University of Bonn, Germany. MH was supported by the Deutsche Forschungsgemeinschaft.

Edinburgh: DJM is supported by an NRS Clinical Fellowship funded by the CSO.

Fran: This research was supported by Foundation FondaMental, Créteil, France and by the Investissements d’Avenir Programs managed by the ANR under references ANR-11-IDEX-0004-02 and ANR-10-COHO-10-01.

Halifax: Halifax data were obtained with support from the Canadian Institutes of Health Research.

iPSYCH BP group: ADB and the iPSYCH team acknowledges funding from The Lundbeck Foundation (grant no R102-A9118 and R155-2014-1724), the Stanley Medical Research Institute, an Advanced Grant from the European Research Council (project no: 294838), and grants from Aarhus University to the iSEQ and CIRRAU centers.

The Mayo Bipolar Disorder Biobank was funded by the Marriot Foundation and the Mayo Clinic Center for Individualized Medicine

Michigan (NIMH/Pritzker Neuropsychiatric Disorders Research Consortium): We thank the participants who donated their time and DNA to make this study possible. We thank members of the NIMH Human Genetics Initiative and the University of Michigan Prechter Bipolar DNA Repository for generously providing phenotype data and DNA samples. Many of the authors are members of the Pritzker Neuropsychiatric Disorders Research Consortium which is supported by the Pritzker Neuropsychiatric Disorders Research Fund L.L.C. A shared intellectual property agreement exists between this philanthropic fund and the University of Michigan, Stanford University, the Weill Medical College of Cornell University, HudsonAlpha Institute of Biotechnology, the Universities of California at Davis, and at Irvine, to encourage the development of appropriate findings for research and clinical applications.

Mount Sinai: This work was funded in part by a NARSAD Young Investigator award to EAS. NeuRA-CASSI-Australia: This work was funded by the NSW Ministry of Health, Office of Health and Medical Research. CSW was a recipient of National Health and Medical Research Council (Australia) Fellowships (#1117079, #1021970).

NeuRA-IGP-Australia: MJG was supported by a NHMRC Career Development Fellowship (1061875).

Norway: TE was funded by The South-East Norway Regional Health Authority (#2015-078) and a research grant from Mrs. Throne-Holst.

Span2: CSM is a recipient of a Sara Borrell contract (CD15/00199) and a mobility grant (MV16/00039) from the Instituto de Salud Carlos III, Ministerio de Economía, Industria y Competitividad, Spain. MR is a recipient of a Miguel de Servet contract (CP09/00119 and CPII15/00023) from the Instituto de Salud Carlos III, Ministerio de Economía, Industria y Competitividad, Spain. This investigation was supported by Instituto de Salud Carlos III (PI12/01139, PI14/01700, PI15/01789, PI16/01505), and cofinanced by the European Regional Development Fund (ERDF), Agència de Gestió d’Ajuts Universitaris i de Recerca-AGAUR, Generalitat de Catalunya (2014SGR1357), Departament de Salut, Generalitat de Catalunya, Spain, and a NARSAD Young Investigator Grant from the Brain & Behavior Research Foundation. This project has also received funding from the European Union’s Horizon 2020 Research and Innovation Programme under the grant agreements No 667302 and 643051.

SWEBIC: We are deeply grateful for the participation of all subjects contributing to this research, and to the collection team that worked to recruit them. We also wish to thank the Swedish National Quality Register for Bipolar Disorders: BipoläR. Funding support was provided by the Stanley Center for Psychiatric Research, Broad Institute from a grant from Stanley Medical Research Institute, the Swedish Research Council, and the NIMH.

Sweden: This work was funded by the Swedish Research Council (M. Schalling, C. Lavebratt), the Stockholm County Council (M. Schalling, C. Lavebratt, L. Backlund, L. Frisén, U. Ösby) and the Söderström Foundation (L. Backlund).

UK - BDRN: BDRN would like to acknowledge funding from the Wellcome Trust and Stanley Medical Research Institute, and especially the research participants who continue to give their time to participate in our research.

UNIBO / University of Barcelona, Hospital Clinic, IDIBAPS, CIBERSAM: EV thanks the support of the Spanish Ministry of Economy and Competitiveness (PI15/00283) integrated into the Plan Nacional de I+D+I y cofinanciado por el ISCIII-Subdirección General de Evaluación y el Fondo Europeo de Desarrollo Regional (FEDER); CIBERSAM; and the Comissionat per a Universitats i Recerca del DIUE de la Generalitat de Catalunya to the Bipolar Disorders Group (2014 SGR 398).

WTCCC: The principal funder of this project was the Wellcome Trust. For the 1958 Birth Cohort, venous blood collection was funded by the UK Medical Research Council.

AHY is funded by the National Institute for Health Research (NIHR) Biomedical Research Centre at South London and Maudsley NHS Foundation Trust and King’s College London. The views expressed are those of the authors and not necessarily those of the NHS, the NIHR, or the Department of Health.

